# (p)ppGpp is required for Virulence of *Shigella flexneri*

**DOI:** 10.1101/2023.08.24.554660

**Authors:** Grace Kago, Charles L. Turnbough, Shelley M. Payne

**Affiliations:** Department of Molecular Biosciences, University of Texas at Austin, Austin, TX, USA; Department of Microbiology, University of Alabama at Birmingham, Birmingham, AL, USA; John Ring LaMontagne Center for Infectious Disease, The University of Texas at Austin, Austin, TX, USA

## Abstract

Infection by the enteric pathogen *Shigella flexneri* requires transit through the gastrointestinal tract and invasion of and replication within the cells of the host colonic epithelium. This process exposes the pathogen to a range of diverse microenvironments. Further, the unique composition and physical environment of the eukaryotic cell cytosol represents a stressful environment for *S. flexneri*, and extensive physiological adaptations are needed for the bacterium to thrive. In this work, we show that disrupting synthesis of the stringent response alarmone (p)ppGpp in *S. flexneri* diminished expression of key virulence genes, including *ipaA, ipaB, ipaC* and *icsA*, and it reduced bacterial invasion and intercellular spread. Deletion of the (p)ppGpp synthase gene *relA* alone had no effect on *S. flexneri* virulence, but disruption of both *relA* and the (p)ppGpp synthase/hydrolase gene *spoT* resulted in loss of (p)ppGpp synthesis and virulence. While the *relA spoT* deletion mutant was able to invade a cultured human epithelial cell monolayer, albeit at reduced levels, it was unable to maintain the infection and spread to adjacent cells, as indicated by loss of plaque formation. Complementation with *spoT* on a plasmid vector restored plaque formation. Thus, SpoT alone is sufficient to provide the necessary level of (p)ppGpp for virulence. These results indicate that (p)ppGpp is required for *S. flexneri* virulence and adaptation to the intracellular environment, adding to the repertoire of signaling pathways that affect *Shigella* pathogenesis.

## INTRODUCTION

*Shigella spp.* are an enteric, pathogenic bacterial genus that is responsible for diarrhea-associated morbidity and mortality, primarily in people living in low and middle-income countries (Kotloff et al., 2013). *Shigella* infections are estimated to cause 80-165 million cases of disease and 600,000 deaths annually (Nemhauser, 2023). *Shigella* are spread via the fecal-oral route in humans (Wharton et al., 1990), and they invade intestinal epithelial cells leading to destruction of the colonic mucosa (Labrec et al., 1964). Of the four characterized *Shigella* species, *Shigella flexneri* is the species associated with endemic bacillary dysentery and is the species most widely used for genetic and virulence studies. The ability of *Shigella spp.* to cause disease requires virulence proteins that are encoded on chromosomal pathogenicity islands (Al-Hasani et al., 2001; Ingersoll et al., 2003; Moss et al., 1999; Vokes et al., 1999) and on a large virulence plasmid (Hale et al., 1983; Kopecko et al., 1980; Sansonetti et al., 1981, 1982). A key virulence component is a Type III secretion system (Blocker et al., 1999) that is essential for injection into the host cell of bacterial effectors that facilitate adhesion, invasion, and evasion of the host innate immune system (Blocker et al., 1999; Brotcke Zumsteg et al., 2014; Konradt et al., 2011).

Because successful colonization by *S. flexneri* requires travel through the environmentally diverse gastrointestinal tract of the host, *S. flexneri* must adapt to multiple chemical and nutritional microenvironments, with differences in pH, O_2_ levels, antimicrobial compounds, and carbon source availability. Upon oral infection, it encounters pH gradients ranging from gastric pH (pH 1-2) to duodenal and ileal pH (pH 6-8), and it traverses the nutrient-rich but antimicrobial mucus layer in the colon (Marteyn et al., 2012; Nuding et al., 2013). Inside the host cell, the unique composition of the eukaryotic cell cytosol requires physiological adaptations for bacteria to survive. Adaptation to the multiple changes in environment requires extensive transcriptional modulation of genes; a gene expression profile study of *S. flexneri* post infection reported either up- or down-regulation of 929 genes in human epithelial HeLa cells and 1060 genes in human macrophage-like U937 cells (Lucchini et al., 2005). Similarly, an analysis of the proteome of intracellular compared to extracellular *Shigella* revealed changes in protein levels corresponding to carbon metabolism enzymes, iron transporters, and outer membrane proteins, among others (Pieper et al., 2013). Additionally, the expression of virulence proteins themselves are subject to regulation by environmental factors. For example, at low temperature and low osmolarity, the transcriptional silencer H-NS inhibits the expression of *virF*, which codes for a key transcription regulator (Falconi et al., 2001; Hromockyj et al., 1992; Porter & Dorman, 1994). VirF regulates *icsA* (Tran et al., 2011), which encodes a surface protein essential for cell-to-cell spread, and *virB,* which encodes a transcriptional regulator of additional virulence genes (Adler et al., 1989). In acidic pH conditions, the CpxA/CpxR two component system also modulates the expression of *virF* (Nakayama & Watanabe, 1995). In anoxic and microaerophilic environments, the transcription factors ArcA and FNR are required for *S. flexneri* invasion and plaque formation (Boulette & Payne, 2007; Marteyn et al., 2010). *S. flexneri* also regulates iron transporters in concert with oxygen availability via FNR and ArcA (Boulette & Payne, 2007).

Adaptation of *S. flexneri* to the intracellular environment of host cells requires responding to the distinct nutritional environment of the eukaryotic cytosol. The cytosol is not a nutrient-rich niche for bacterial growth; extracellular or vacuolar pathogens microinjected into the eukaryotic cytosol were unable to grow (Goetz et al., 2001). Moreover, studies in *Legionella pneumophila*, *Anaplasma phagocytophilum*, and *Francisella tularensis* indicated that the basal concentrations of amino acids in the eukaryotic cell are insufficient for bacterial proliferation, leading to the idea of amino acids as “nutritional rheostats” for infectious bacteria (Abu Kwaik & Bumann, 2013). Another aspect of what has been termed “nutritional immunity” is the limited availability of iron in the host (Skaar, 2010; Sousa Gerós et al., 2020; Weinberg, 1977). Iron is restricted in the host cell cytoplasm (Payne et al., 2006; Sousa Gerós et al., 2020), and the ability of *S. flexneri* to replicate in host cells is dependent on their expression of high-affinity iron transport systems, in particular the ferrous iron transport Sit (Fisher et al., 2009; Runyen-Janecky et al., 2003).

The nutrient limitations initially encountered by *S. flexneri* upon entering the host cell cytosol trigger signaling pathways that allow *Shigella* to modulate gene expression and adapt to this environment. For example, *S. flexneri* alter their gene expression upon entry into the host cells to metabolize pyruvate, and they use the mixed acid fermentation pathway when growing intracellularly (Kentner et al., 2014; Pieper et al., 2013; Waligora et al., 2014). Formate is produced as a by-product of mixed-acid fermentation, and the secreted, extracellular formate serves as a signal to regulate *S. flexneri* virulence gene expression inside the cytosol of infected host cells. However, the specific signaling pathways and bacterial intracellular signals that control these responses during infection are not fully characterized. (Koestler et al., 2018).

A primary adaptive response to nutrient deficit in many bacterial species is the stringent response. The bacterial stringent response is mediated by the hyperphosphorylated nucleotides guanosine tetraphosphate (ppGpp) and guanosine pentaphosphate (pppGpp) (Cashel & Gallant, 1969) collectively referred to as (p)ppGpp. Intracellular concentrations of (p)ppGpp increase from micromolar to millimolar levels upon nutrient imitation (Kuroda et al., 1997; Steinchen et al., 2018; Varik et al., 2017), and they are synthesized by proteins which belong to the RelA/SpoT Homologue (RSH) family (Atkinson et al., 2011; Gentry & Cashel, 1996; Mittenhuber, 2000).

In many species of β- and γ-proteobacteria, the ancestral gene that encodes the RSH protein has been duplicated into two genes, *relA* and *spoT* (Mittenhuber, 2000). The ribosome-associated (p)ppGpp synthase RelA is activated upon amino acid depletion, specifically by the presence of uncharged tRNAs in the ribosome A site (Arenz et al., 2016; Kushwaha et al., 2019; Winther et al., 2018). In contrast, SpoT, a (p)ppGpp synthase and hydrolase (An et al., 1979; Laffler & Gallant, 1974) maintains a basal level of (p)ppGpp in the cell in the absence of nutritional stress (Fernández-Coll & Cashel, 2020). SpoT is activated by deficiencies in fatty acids (Battesti & Bouveret, 2006), phosphate (Rao et al., 1998; Spira et al., 1995), nitrogen(Brown et al., 2014) carbon (Meyer et al., 2021), and iron (Vinella et al., 2005).

(p)ppGpp levels modulate key cellular processes including transcription, DNA replication, translation, (Gourse et al., 2018; Srivatsan & Wang, 2008), GTP synthesis (Kriel et al., 2012), and ribosome assembly (Corrigan et al., 2016). Even at low concentrations, (p)ppGpp has been shown to influence the transcription of hundreds of genes, with (p)ppGpp regulation important for cell fitness even in the absence of known stressors (Kundra et al., 2020).

In addition to critical roles in cellular metabolism, (p)ppGpp modulates a host of virulence phenotypes including adhesion, biofilm formation, toxin production, motility, and antibiotic resistance and tolerance (reviewed in: Chau et al., 2021; Dalebroux et al., 2010; Gaca et al., 2013). In pathogens that invade eukaryotic cells, (p)ppGpp has been reported to be essential for several attributes of virulence. For example, in *Salmonella enterica*, a mutant strain without (p)ppGpp had decreased expression of the virulence plasmid and the transcriptional activators *hilA and invF*, which are important for *Salmonella* virulence (Pizarro-Cerdá & Tedin, 2004). (p)ppGpp was also shown to be essential for both adherence and activation of transcription regulators in enterohemorrhagic *Escherichia coli* (Nakanishi et al., 2006). A *Campylobacter jejuni* ppGpp-null (ppGpp^0^) strain was reported to have defects in adherence, invasion, and intracellular survival (Gaynor et al., 2005). Similarly, a (p)ppGpp^0^ strain of *Brucella abortus* showed reduced invasion of mouse spleens (Kim, 2005). Gram positive pathogens also require (p)ppGpp for key steps in virulence. In *Streptococcus suis,* adherence to and invasion of Hep-2 cells was decreased in strains without (p)ppGpp (Zhu et al., 2016), and in *Enteroccoccus faecalis*, strains with reduced amounts of (p)ppGpp showed diminished invasion of coronary artery endothelial cells (Colomer-Winter et al., 2018).

In this study, we investigated the role of (p)ppGpp synthesis in *Shigella* virulence. We generated a (p)ppGpp^0^ strain of *S. flexneri* by deleting both *relA* and *spoT*. The (p)ppGpp^0^ strain demonstrated an invasion defect, lower amounts of virulence proteins, intracellular filamentation, and reduced localization of the polar autotransporter protein IcsA resulting in an inability to spread and maintain infection.

## RESULTS

### *relA* and *spoT* are conserved across *Shigella spp*

Genomic alignments of *relA* and *spoT* from the genomes of *S. flexneri, Shigella sonnei, Shigella dysenteriae,* and *Shigella boydii*, as well as the enterohemorrhagic *E.* coli O157:H7 showed conservation of the genes across these species (Fig.1A), suggesting that *Shigella spp.,* like other enteric bacteria produce (p)ppGpp as a signaling molecule. We used an *in vitro* reporter to measure (p)ppGpp in a wild**-**type *S. flexneri* strain compared to the isogenic *ΔrelA ΔspoT* mutant. To generate this reporter system, a plasmid was constructed that carried the (p)ppGpp-activated *csgD* promoter of *E. coli* (Sanchez-Vazquez et al., 2019) inserted upstream of a promoterless *E. coli lacZ gene* (Farinha & Kropinski, 1990). Comparison of reporter-carrying wild**-**type, *ΔrelA* (ΔR), and *ΔrelA ΔspoT* (ΔRS), strains of *S. flexneri* on medium containing the β-galactosidase indicator X-gal showed similar enzyme activity (i.e., blue color) for the wild-type and ΔR *strains* (Fig. 1B). In contrast, the ΔRS colonies were colorless, with β-galactosidase only detected in the densest area of the streak, demonstrating the (p)ppGpp deficiency of the double mutant (Fig. 1B). We also measured *pcsgD-lacZ* expression (i.e., b-galactosidase levels) in mid-log and stationery-phase cells of the wild-type DR, and DRS strains grown in liquid medium (Fig. 1C). The results show greatly reduced reporter gene expression in the DRS strain compared to the other two strains, consistent with a lack of (p)ppGpp in the DRS strain.

**Figure 1.**
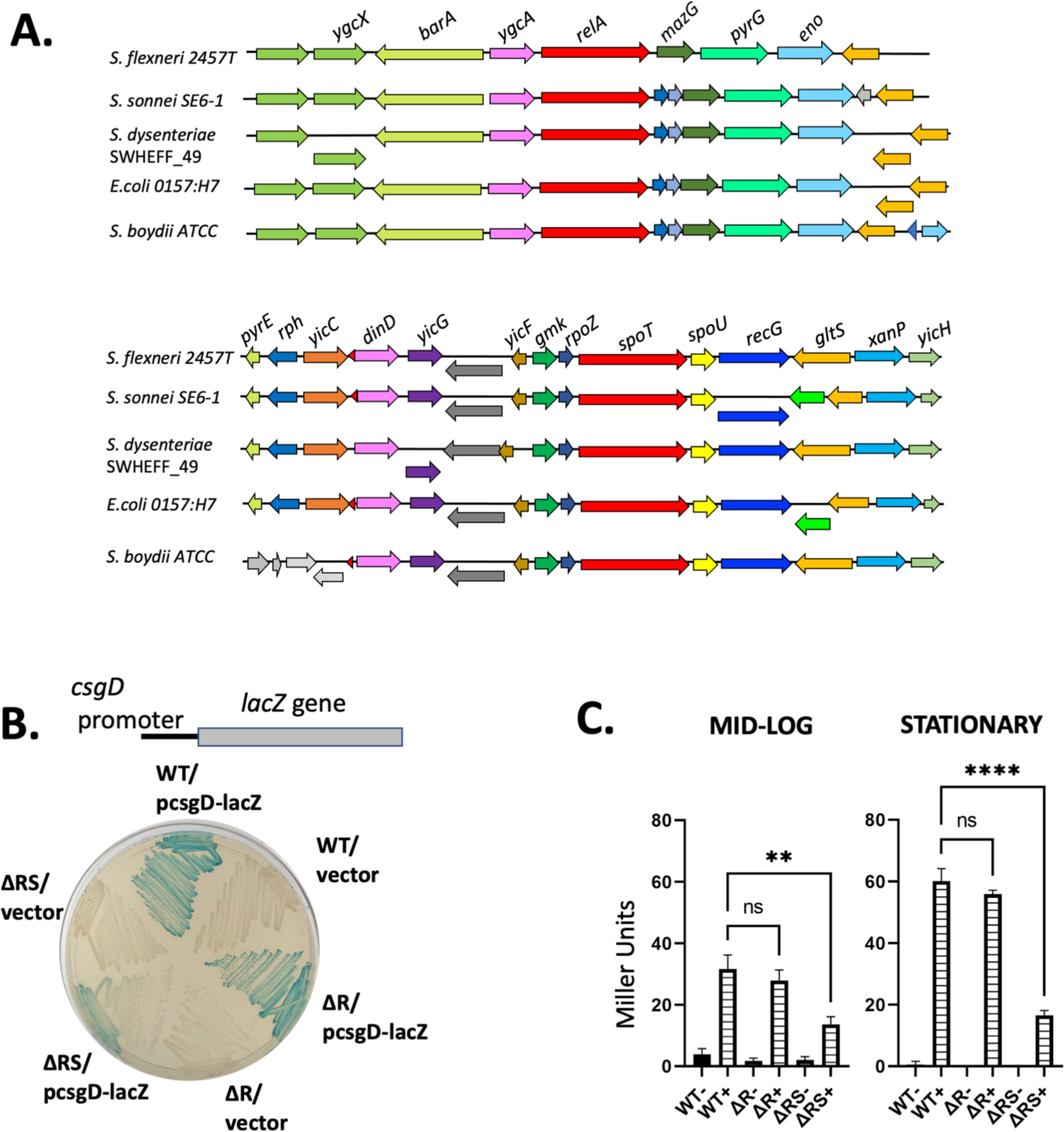
*relA* and *spoT* are present in *Shigella spp.* and SpoT is required for efficient plaquing. A) Genomic alignments of *relA* and *spoT* from the genomes of *S. flexneri, S. boydii, S. sonnei*, *S. dysenteriae,* and the enterohemorrhagic *E. coli* O157:H7. Genomes were sourced from NCBI database (NCBI access numbers listed in Methods). Alignments and annotations were conducted with RAST. Arrows below the lines indicate genes that overlap adjacent open reading frames. B) *S. flexneri* strains carrying the *csgD* promoter fused to *lacZ* in pQF50 (*pcsgd-lacZ*) or the pQF50 alone (*vector*) were streaked on L agar with X-gal. The wild-type (WT) and ΔR strains, which retain a functional SpoT, produced β-galactosidase, as indicated by the blue colonies, while the ΔRS strain fails to express *lacZ*. C) Miller assays to quantitate β-galactosidase activity in the strains reported in (B) grown to mid-log and stationary phase. (**) indicates p value <0.01, (****) p value <0.0001, and not significant (ns) via ordinary one-way Analysis of Variance (ANOVA).

### SpoT is necessary for efficient invasion and intercellular spread of *S. flexneri*

To determine whether (p)ppGpp plays a specific role in *S. flexneri* virulence, wild-type, ΔR, and ΔRS strains were compared in a plaque assay on Henle cells. The plaque assay tests for characteristics of wild-type *S. flexneri* that include (i) invasion (ii) replication and (iii) intercellular spread to adjacent cells which produces clear plaques in the Henle cell monolayer (Oaks et al., 1985). Infection of the monolayer by the ΔR single mutant strain resulted in wild-type size plaques, whereas the double mutant (ΔRS) was unable to form plaques (Fig. 2). A mutant with only *spoT* deleted could not be tested, because (p)ppGpp hydrolysis by SpoT is essential to ameliorate the toxic effects of (p)ppGpp overproduction and accumulation, and a strain with disruption of the *spoT* gene in the presence of an intact *relA* gene is not viable (Xiao et al., 1991). Complementation of the double mutant with a plasmid carrying the wild-type *spoT* gene (pWKS30-spoT) resulted in restoration of plaque formation (Fig. 2). Thus, *spoT* produces sufficient (p)ppGpp in response to the intracellular environment for *S. flexneri* invasion, replication, and spread.

**Figure 2.**
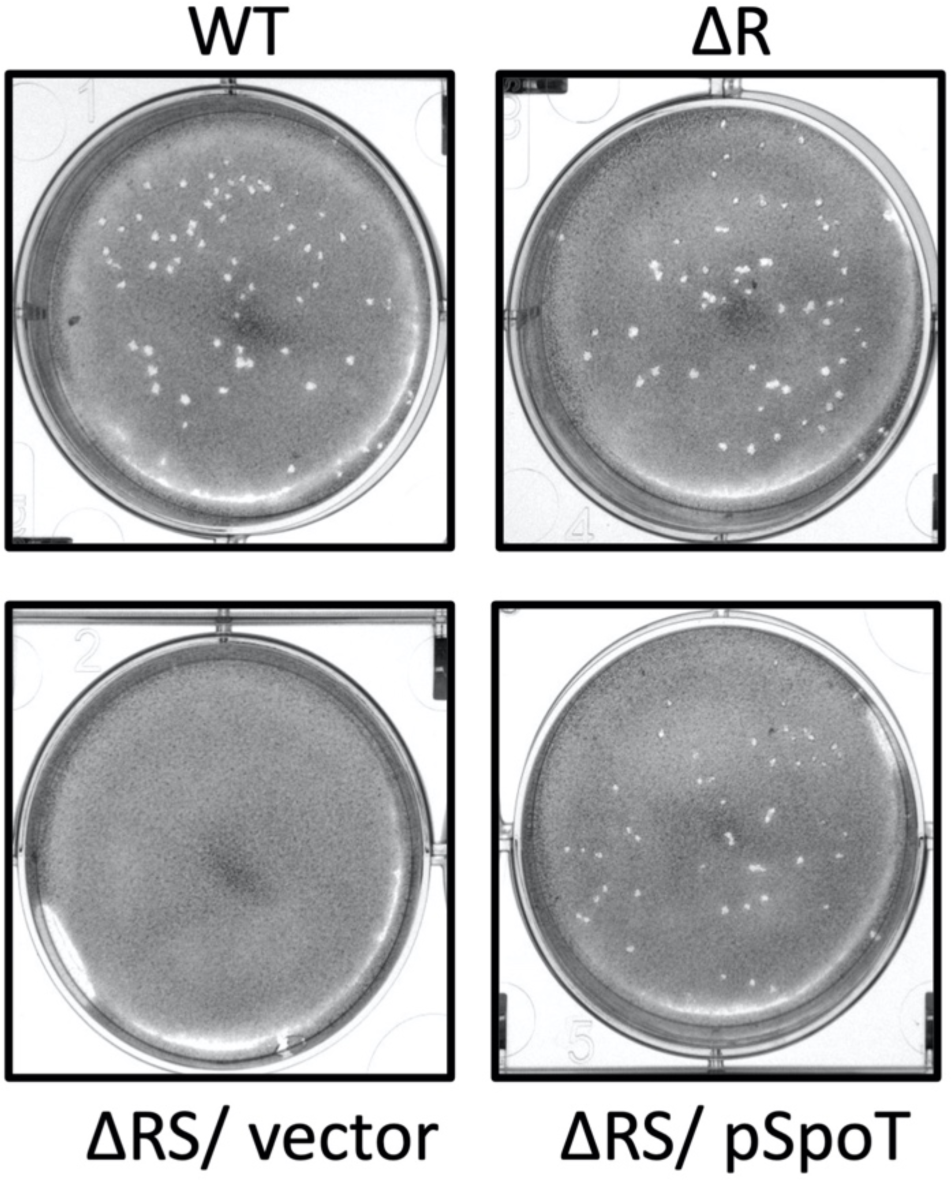
SpoT is necessary for *S. flexneri* plaque formation. Plaque assay of WT, βR, and the βRS strains with (βRS/ pSpoT) and without (ΔRS/ vector) cells complementation of *spoT*.

### The SpoT mediated virulence defect is overcome by elevated Hfq

In proteobacteria, (p)ppGpp directly affects transcription by binding to two sites on RNA polymerase (RNAP); the first at the interface of the β′ and ω subunits and the second at the interface of the β′ secondary channel and the transcription factor DksA (Gourse et al., 2018; Myers et al., 2020; Sanchez-Vazquez et al., 2019). DksA is necessary for both positive and negative transcriptional regulation in response to high concentrations of (p)ppGpp (Fernández-Coll et al., 2020; Paul et al., 2004, 2005; Sanchez-Vazquez et al., 2019). One of the genes positively regulated by DksA and (p)ppGpp encodes the pleiotropic regulator protein Hfq, an RNA chaperone (review: (Chao & Vogel, 2010; Sharma & Payne, 2006). Hfq is necessary for virulence in *S. flexneri*; an *hfq* mutant fails to form plaques, and the same phenotype is observed with a *dksA* mutant (Sharma & Payne, 2006). Although DksA is expected to affect transcription of a large number of genes in addition to *hfq* in *S. flexneri*, the primary role of DksA in *Shigella* virulence is its requirement for *hfq* expression; expressing *hfq* independently of DksA restored plaque formation in the *dksA* mutant (Sharma & Payne, 2006). If the observed virulence defect of the ΔRS strain was caused by the lack of (p)ppGpp needed to induce Hfq synthesis, expressing *hfq* independently of (p)ppGpp/DksA should bypass the need for (p)ppGpp and restore plaque formation. We introduced a plasmid with an inducible *hfq* and determined plaque formation with increasing expression of *hfq* (Fig. 3B). In the presence of increasing amounts of inducer, the level of Hfq increased in both the wild-type and the ΔRS strains, as shown by Western blot analysis (Fig. 3C), but this did not restore plaque formation by the mutant (Fig. 3B). This result indicates that the effect of (p)ppGpp on Hfq production is not the sole reason for reduced virulence of the ΔRS mutant.

**Figure 3.**
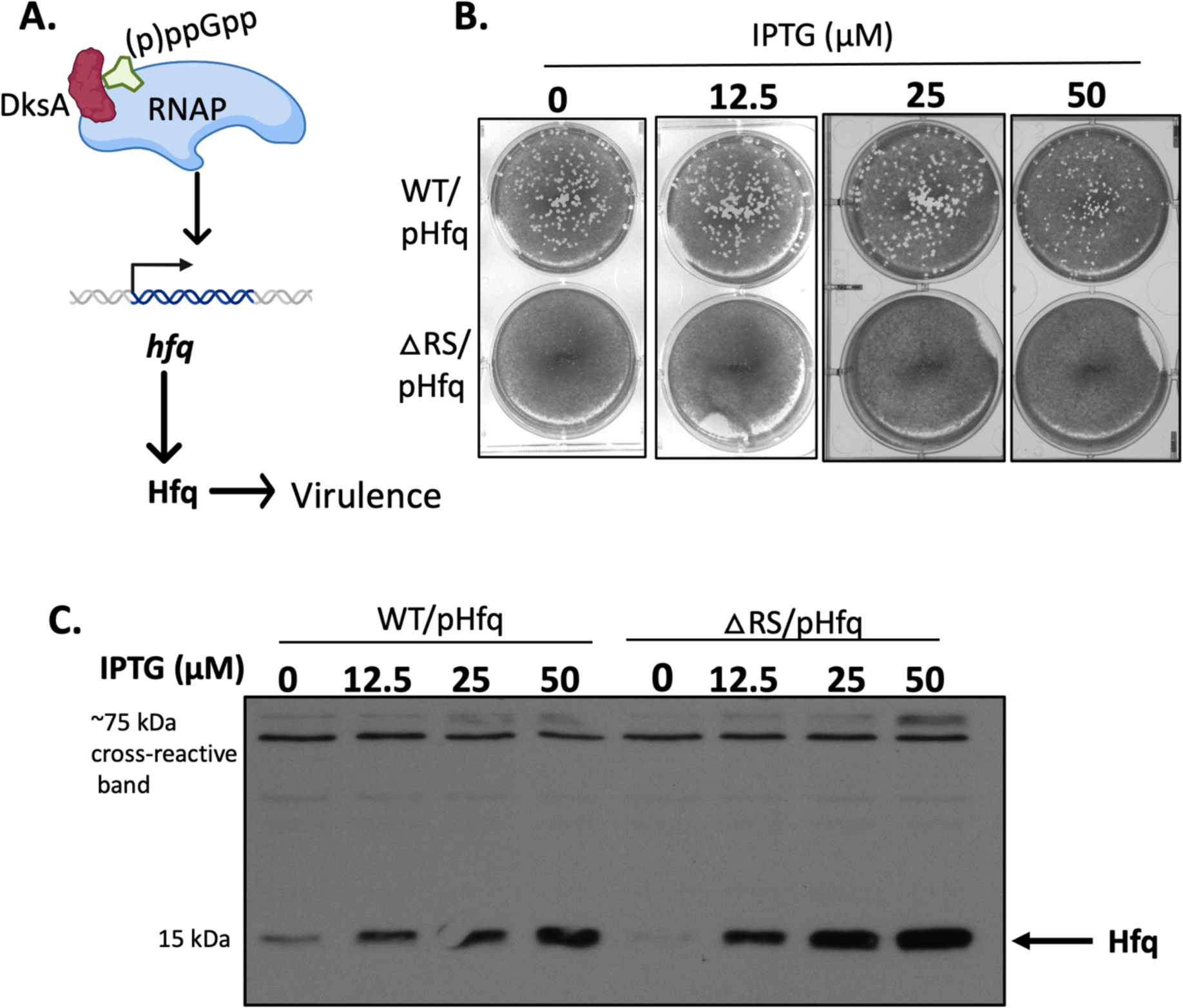
The SpoT mediated plaque defect is independent of Hfq. A) Schematic of the relationship between (p)ppGpp, DksA, Hfq and *S. flexneri* virulence. B) Image of plaque assay conducted using WT and βRS strains of *S. flexneri* carrying the IPTG-inducible *hfq* expression plasmid (Sharma & Payne, 2006). *S. flexneri* strains were grown with the indicated amounts of IPTG, and that concentration was maintained throughout the infection and plaque assay. Plates were stained with crystal violet. C) Immunoblot indicating Hfq expression as a function of IPTG concentration was conducted using polyclonal rabbit anti-Hfq antibody. A 75-kDa cross-reactive band used as a loading control. As noted previously (Sharma & Payne, 2006), high levels of Hfq (e.g. induction with 50 µM IPTG) reduced the size and number of the plaques in the wild-type strain (Sharma & Payne, 2006).

### SpoT is necessary for invasion and spread

Invasion assays showed that the ΔRS strain, but not the ΔR strain, had a partial defect in invading Henle cells (Figs. 4A and B). To determine whether SpoT is only needed for invasion or is impacting virulence at subsequent stages, ΔRS expressing SpoT under an arabinose inducible promoter (pBAD24-SpoT) was used to identify the time at which SpoT is required. The strains were grown in the absence of arabinose and allowed to invade a Henle cell monolayer in the absence of arabinose. Arabinose was then added post-invasion to monolayers infected with the strain containing the empty vector or the inducible SpoT vector. In the absence of arabinose, cells infected with the ΔRS strain harboring pBAD24-SpoT made no plaques. However, plaques were observed in the presence of arabinose (Fig. 4C), the same phenotype as the ΔRS cells harboring *spoT* expressed under a constitutive promoter on plasmid pWKS30-SpoT (Fig. 4C). This result indicates that there is sufficient virulence gene expression in the absence of SpoT for *S. flexneri* to invade Henle cells, but *spoT* expression, (i.e. (p)ppGpp production) is necessary for *S. flexneri* to maintain the infection and spread cell-to-cell to form plaques (Fig. 4C).

**Figure 4.**
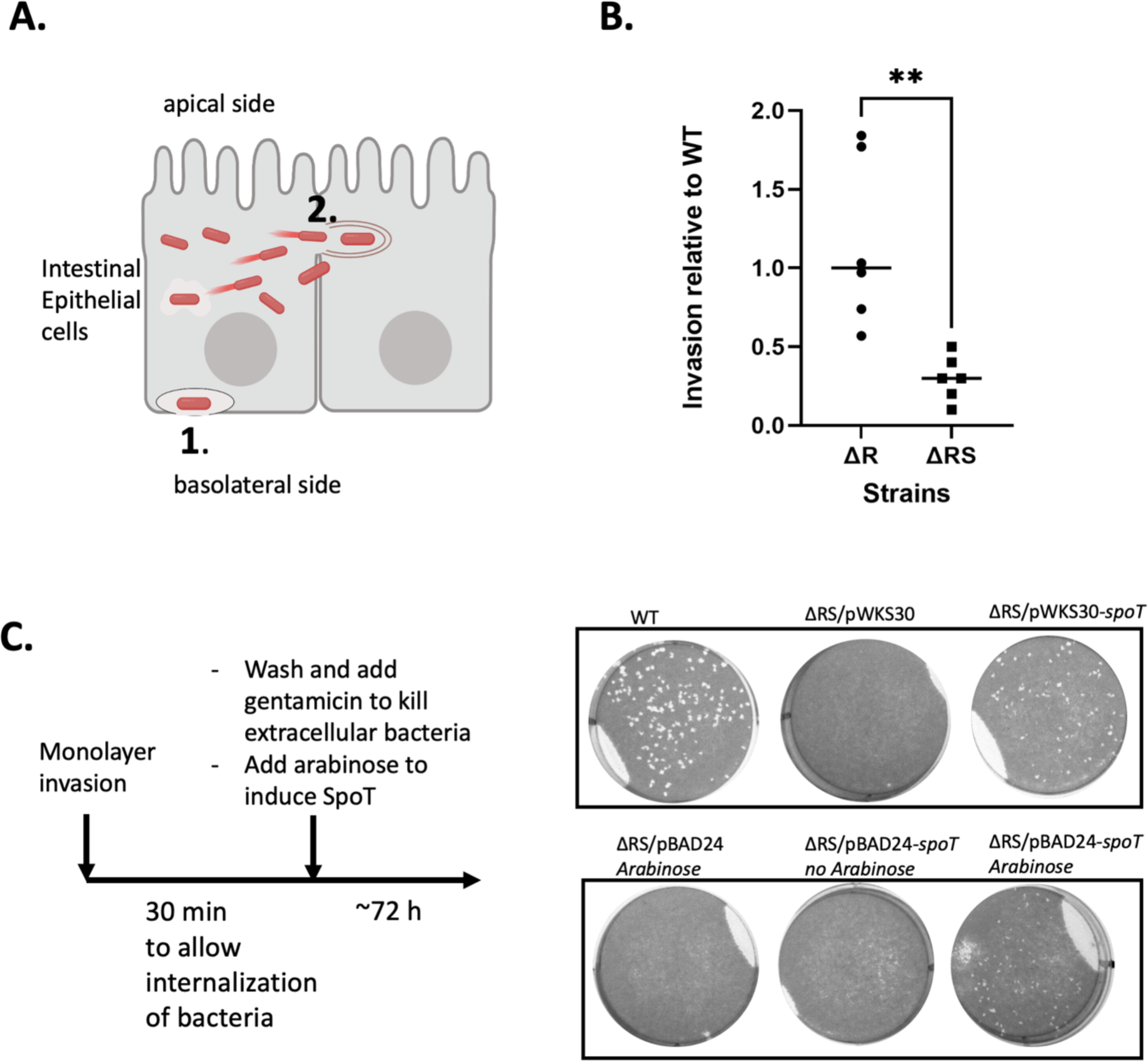
Invasion is reduced in the ΔRS strain, and SpoT is necessary and sufficient for intercellular spread. A) Schematic showing steps in *S. flexneri* virulence, (1) invasion and (2) spread to adjacent epithelial cells. B) Invasion of *S. flexneri* ΔR and ΔRS strains relative to wild-type (WT), (**) p<0.05, Students *t* test. C) Plaque assay of WT and ΔRS strains and the effect of constitutive or induced *spoT* expression. Schematic at left shows the timing of induction of *spoT* from the arabinose promoter on pBAD24. The Henle cells were infected with the wild-type (WT) *S. flexneri* or with the ΔRS mutant containing constitutively expressed *spoT* cloned into pWKS30 (ΔRS/pWKS30-*spoT*) or in the arabinose-inducible vector pBAD24 (ΔRS/pBAD24-*spoT*). The ΔRS mutant containing each of the empty vectors (ΔRS/pWKS30 and ΔRS/pBAD24) were included as negative controls. Henle cells were infected with strains grown in the absence of arabinose, allowed to invade. Extracellular bacteria were removed by washing, and gentamicin was added to kill any remaining extracellular bacteria prior to addition of arabinose. Schematic created with Biorender.com

### SpoT is necessary for IcsA localization

*S. flexneri* are non-motile, and they rely on actin-based movement for intercellular spread. Actin based motility requires the expression and efficient polar localization of the autotransporter protein IcsA (Goldberg & Theriot, 1995). Because the ΔRS mutant failed to form plaques, despite retaining moderate invasion of the cells, a defect in intercellular motility is consistent with this phenotype. Therefore, indirect immunofluorescence was used to assess IcsA localization. Compared to the wild-type strain, the ΔRS strain showed diminished polar localization of IcsA both *in vitro* (Fig. 5A) and inside Henle cells (Fig. 5B). This result indicates that (p)ppGpp is required for *icsA* expression, IcsA synthesis, and/or polar localization of IcsA.

**Figure 5.**
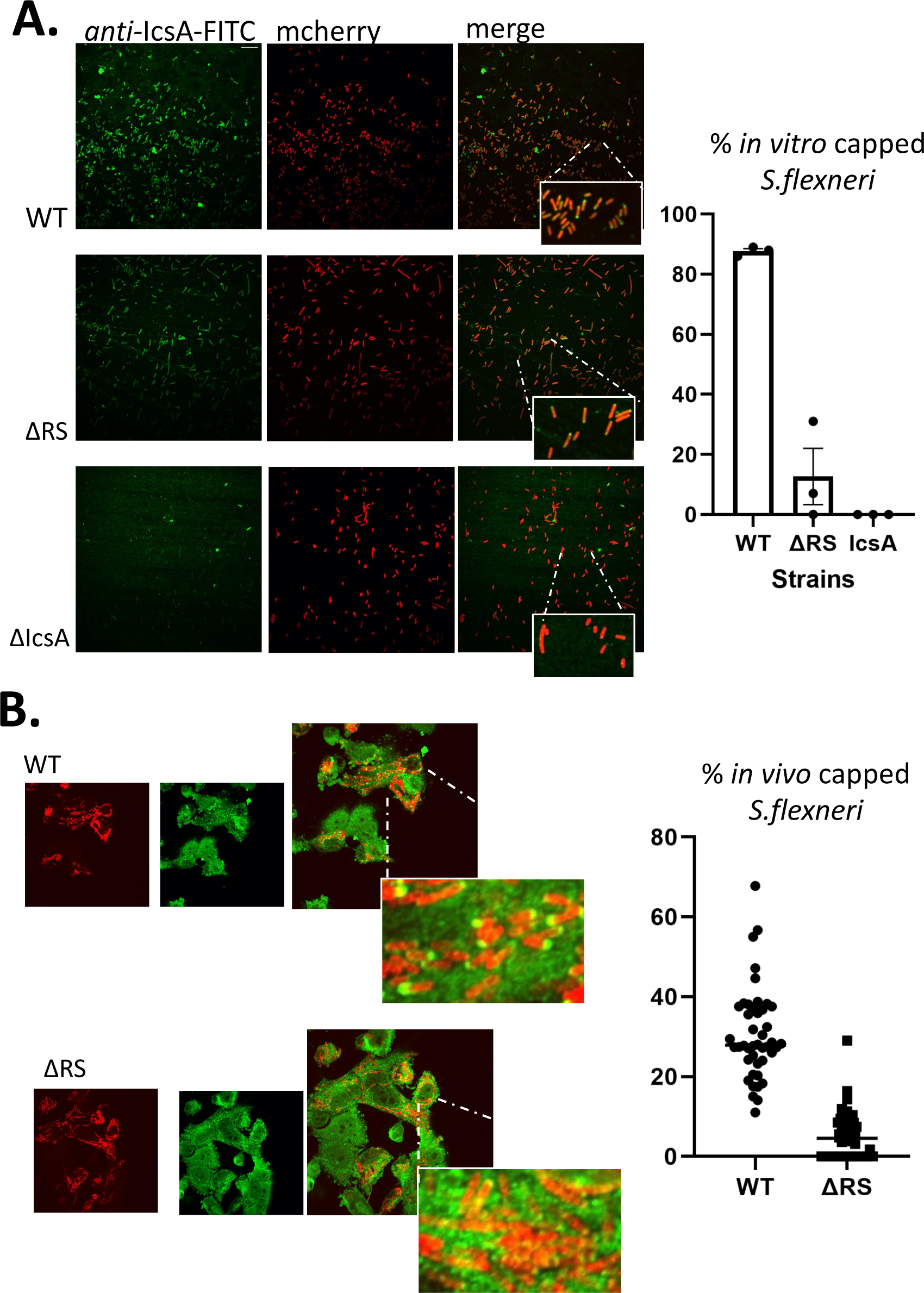
SpoT is necessary for IcsA localization. A) The average percentage of capped cells in the WT and ΔRS strains grown *in vitro* was determined via indirect immunofluorescence of IcsA using anti-IcsA antibody. Data represents three biological replicates, and 100 cells from each strain were counted for each replicate. Representative images are shown. B) *In vivo* indirect immunofluorescence of surface IcsA of WT and ΔRS strains after infection of Henle cells. *S. flexneri* strains expressing m-Cherry were stained with anti-IcsA antibody; Each dot represents the % of *S. flexneri* with IcsA at one pole per image. Forty-five images containing a total of 5112 cells were counted for the wild-type strain, and 50 images with 3137 cells of the ΔRS strain were observed. Data were from three biological replicates. Representative images are shown.

Images of the intracellular bacteria showed that many of the ΔRS cells were much longer than cells of the wild-type strain (Fig. 5B). The abnormal morphology could affect polar localization of IcsA. To assess the effect of cell elongation on IcsA localization, the bacteria were exposed to a sublethal concentration of cephalexin to induce elongation (Chung et al., 2011) and stained with anti-IcsA antibody. The elongated cells were still able to localize IcsA to the pole (Fig. 6), indicating that elongation *per se* is not responsible for the loss of IcsA caps.

**Figure 6:**
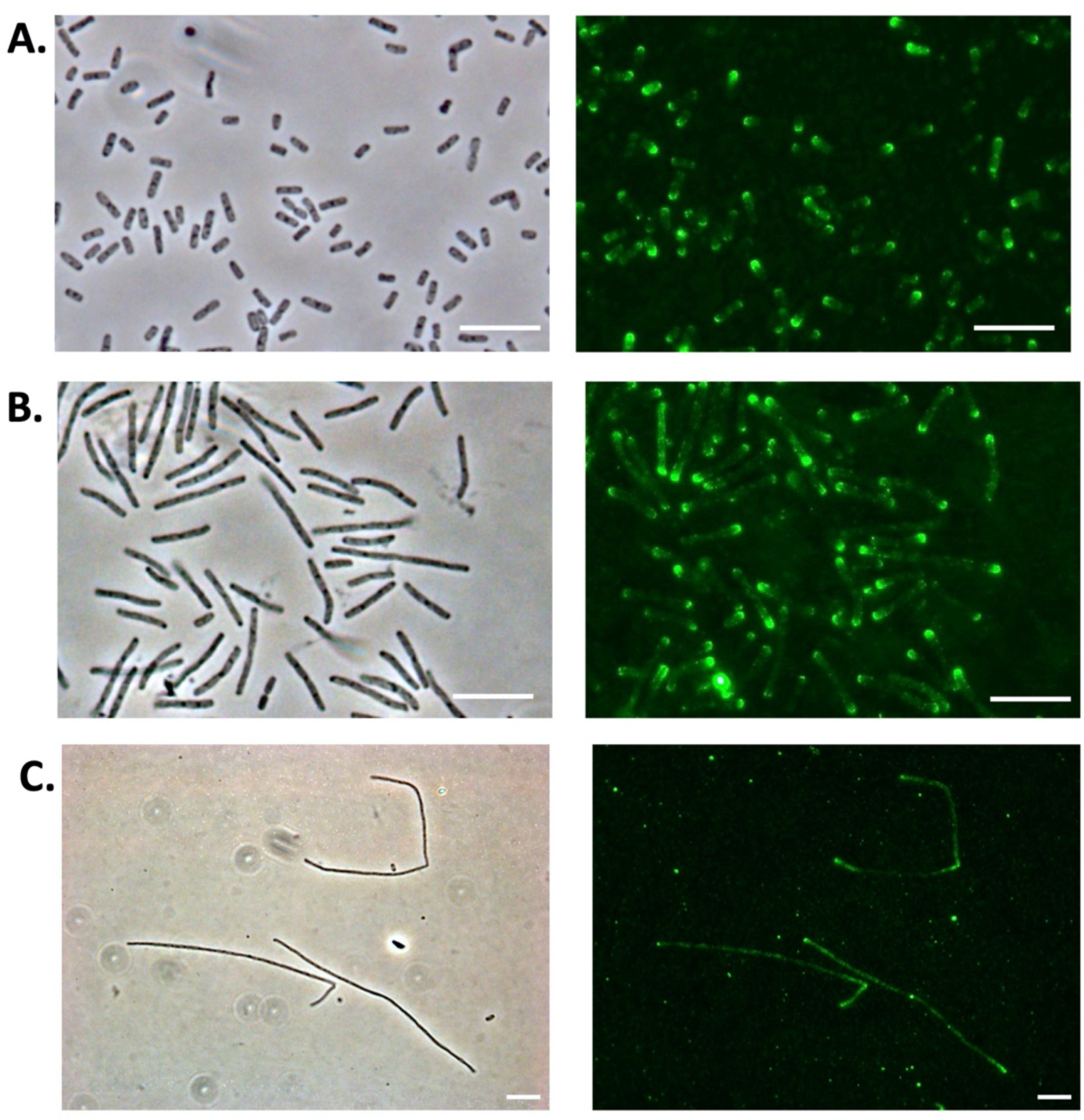
Filamentation does not disrupt polar localization of IcsA. Wild-type *S. flexneri* were grown in LB with or without cephalexin, stained with antibody against IcsA, and imagined using phase contrast (left) and indirect immunofluorescence (right). **A)** cells grown without cephalexin **B)** cells grown with cephalexin for 1hr, and **C)** cells grown with cephalexin for 2 h and 45 min and then stained. Scale bar indicates 5 microns.

### SpoT modulates virulence protein expression

To assess the levels of IcsA and other virulence proteins in the ΔRS strain, Western blot analysis with monkey convalescent (post *S. flexneri* infection) antiserum was used. Bands corresponding to the virulence proteins IpaA, IpaB, IpaC and IcsA were lower in intensity with the ΔRS mutant than with the wild-type strain, indicating a lower amount of the virulence proteins (Fig. 7A). This deficit was restored for the most part upon complementation with *spoT* (Fig. 7A). The *S. flexneri* strain CFS100, which lacks the virulence plasmid, was used as a negative control to confirm that the protein bands corresponded specifically to *Shigella* virulence proteins (Fig. 7A). Because expression of the *ipa* operon and *icsA* gene is regulated by VirF, we determined the level of VirF in both the wild-type and ΔRS strains. S-tagged VirF encoded on the low copy number vector pWKS30 was introduced into the wild-type and ΔRS strains. The level of VirF was measured via an immunoblot with antiserum against the S-tag epitope. Bands corresponding to both forms of VirF, VirF_30_ and VirF_21_, were observed at higher intensity in the wild type compared to the ΔRS strain (Fig. 7B). The reduced amount of VirF in the mutant indicates that (p)ppGpp has a role in maintaining the levels of VirF necessary for enhanced expression of Ipa proteins and IcsA. The reduction in amount of VirF appears to be post-transcriptional. Measurement of the *virF* mRNA levels showed that there was no difference in the amount of mRNA between the wild-type and ΔRS strains (Fig. 7C).

**Figure 7.**
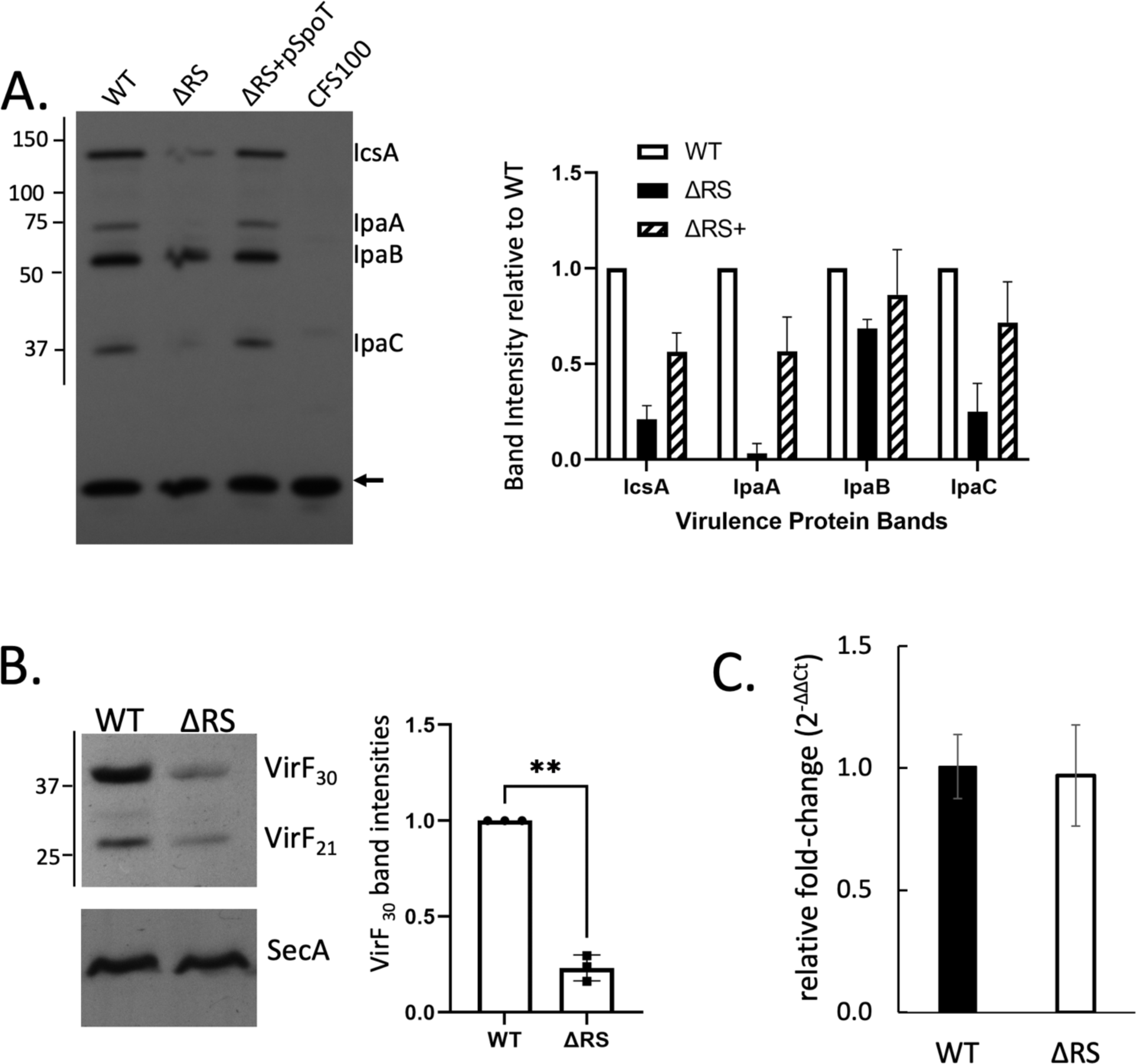
SpoT is necessary for virulence protein expression. A) Immunoblot of *S. flexneri* proteins from the wild type (WT), ΔRS, ΔRS complemented with plasmid-encoded SpoT (ΔRS + pSpoT), and CSF100, the control strain which lacks the virulence plasmid, was probed with monkey convalescent primary antiserum to image virulence proteins. Numbers at left indicate sizes (in kDa) of MW markers. Identification of protein bands is indicated on the right. The arrow indicates an *S. flexneri* protein recognized by the antiserum that is not encoded on the virulence plasmid and is used as a loading control. The immunoblot is representative of three independent experiments. B) Immunoblot to detect S-tagged VirF_30_ and VirF_21_ was probed with anti-S-tag antibody to compare the wild-type and ΔRS mutant. Numbers at left indicate sizes (in kDa) of MW markers. Blot is representative of three biological replicates. The migration of the affinity-tagged recombinant VirF_21_ and VirF_30_ is slightly higher than expected as previously reported (Skovajsová et al., 2022). The upper portion of the blot was cut off and probed with anti-SecA antibody as a loading control. C) Relative *virF* expression in WT and ΔRS strains of *S. flexneri* grown to log phase was measured by RT-qPCR. *C_T_* values were normalized to the mean for an endogenous control, *secA*, and relative fold change (2^-ΔΔCt^) reported. Measurements were derived from three biological replicates, and error bars represent the standard deviation of the mean.

### Elongation of ΔRS cells during intracellular growth

Elongation of cells of the ΔRS strain in the Henle cell cytoplasm was not the cause of improper IcsA localization, it did indicate that the mutant was unable to control cell length and division in response to the intracellular environment (Fig 8A). Measurement of the length of the wild-type and ΔRS strains in rich medium *in vitro* showed that the mutant was slightly longer on average, with a wider range of cell lengths (Fig. 8B and 8C left-side panels). Filamentation, such as that noted with the intracellular ΔRS cell has been noted in *E. coli* in response to exposure to stressful environments, including peroxide (Imlay & Linn, 1987) and antibiotics (Rolinson, 1980) and during infection (Justice et al., 2006). To determine whether the ΔRS mutant filamented in response to stress, the cells were grown in high osmolarity or with iron limitation. As was seen during intracellular growth (Fig. 5B), growth of the cells in vitro in high salt (340 mM) or in iron restriction (presence of an iron chelator) resulted in cell elongation (Fig. 8B andd 8C, right-side panels). Thus, the elongation of the ΔRS mutant is consistent with (p)ppGpp being required for adaptation to the physical environment within the human cell.

**Figure 8.**
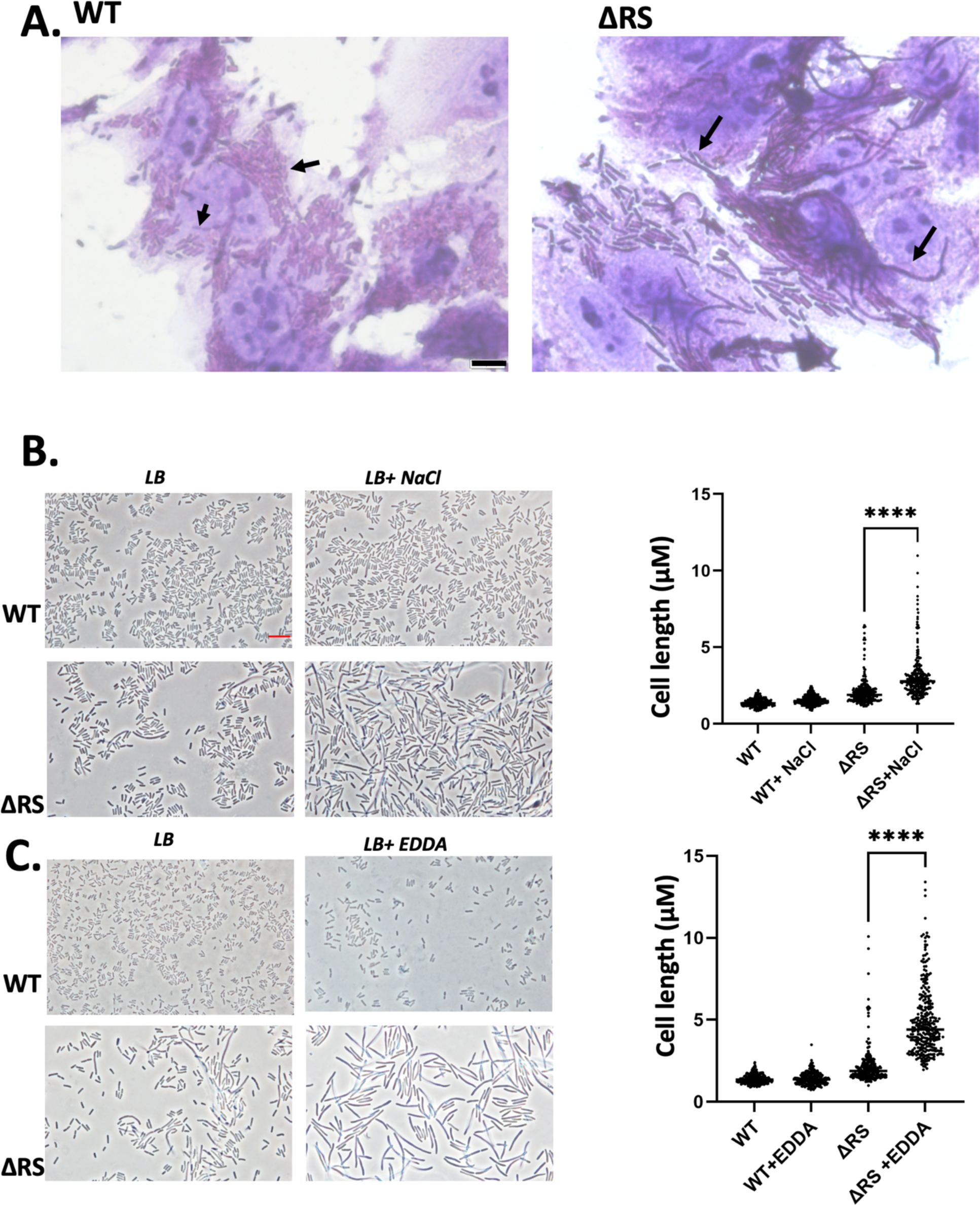
*S. flexneri* forms filaments inside host cells and in response to osmotic stress. A. Henle cells were infected with the wild-type (WT) or ΔRS strains, and the infected cells were stained and imaged 3 h post infection. Arrows indicate intracellular bacteria. B and C. Phase contrast images of cells grown for 3 h in LB with 170 or 340 mM NaCl (B) or with 0 or 250 µg/ml of the iron chelator ethylenediamine di (*ortho*-hydroxyphenylacetic acid) (EDDA) (C). The graphs at right shows the distributions of cell lengths for 100 bacteria per condition per replicate, with three biological replicates of each. (****) indicates statistical significance between lengths of treated vs. untreated ΔRS strain lengths by student’s *t* test p < 0.0001

## DISCUSSION

To cause disease, *S. flexneri* must invade, replicate, and spread intercellularly to adjacent cells. Here we show that SpoT, a protein that is essential for controlling (p)ppGpp concentrations in *Enterobacteriacea*, is necessary for *S. flexneri* virulence. Because (p)ppGpp is a major regulatory signal, or alarmone (Stephens et al., 1975), needed for regulating biosynthetic pathways in response to stress signals, it is not surprising that it would affect the ability of *S. flexneri* to adapt to and survive in the host environment. Here we show that (p)ppGpp is required for efficient expression of the virulence genes that are essential for invasion and cell-to-cell spread.

Two genes, *relA* and *spoT,* control the synthesis of (p)ppGpp. Because of the variable concentration of free amino acids in the eukaryotic cytosol (Piez & Eagle, 1958), and the documented amino acid auxotrophy of Δ*relA* strains (Xiao et al., 1991) we hypothesized that deleting the *relA* gene would be sufficient to test (p)ppGpp’s role in allowing growth of *S. flexneri* in the eukaryotic cytosol. However, the Δ*relA s*train demonstrated no defect in either invasion or spread, whereas deleting *spoT* in the *relA* deletion strain resulted in decreased invasion and no spread. SpoT is responsible for basal (p)ppGpp synthesis and increased (p)ppGpp synthesis under certain stress conditions, indicating that one of these activities is important in modulating *Shigella* virulence.

Although we showed that expression of wild-type levels of multiple virulence proteins required SpoT, and presumably (p)ppGpp synthesis, a major mechanism by which SpoT/(p)ppGpp impacts *Shigella* virulence is via polar localization of the actin polymerization factor IcsA. In the wild-type strain, IcsA forms a cap at one pole of the bacterium, which results in unipolar polymerization of actin and propulsion of the bacterium in the cytosol and into neighboring cells (Goldberg et al., 1993; Goldberg & Theriot, 1995). However, far fewer IcsA caps were present on the ΔRS cells, and this correlated with loss of intercellular spread of the mutant.

The reduced invasion and loss of intercellular spread by the ΔRS mutant may be partially because of the effects of (p)ppGpp that are mediated through DksA. DksA activates expression of Hfq, which is required for *Shigella* virulence (Sharma & Payne, 2006). Ectopic expression of Hfq in the *dksA* mutant restores virulence, but it does not restore virulence of the ΔRS strain. Thus, functions of (p)ppGpp, in addition to its role in Hfq synthesis are important in *Shigella* virulence. These functions may be independent of DksA.

In a previous study to determine which proteins *S. flexneri* synthesized when it was growing in the cytoplasm of host cells (Pieper et al., 2013), we did not observe increased amounts of stress response proteins, but there were changes in proteins responsible for key metabolic pathways including iron transport and carbon metabolism. Because both iron (Vinella et al., 2005) and carbon (Traxler et al., 2006) flux have been implicated in SpoT activation, it is possible that these signals activate SpoT to synthesize (p)ppGpp in intracellular *S. flexneri* and help regulate the levels of the virulence regulator VirF and its downstream virulence genes. *virF* also is responsive to osmolarity (Porter & Dorman, 1994) and other environmental stressors as reviewed by: (Di Martino et al., 2016). *Shigella spp.* likely have evolved to respond to multiple environmental cues to ensure that they appropriately express virulence genes in the epithelial cytosol to promote replication and spread. While much of the known virulence gene regulation is at the transcriptional level, the modulation of VirF protein levels in response to (p)ppGpp appears to be post-transcriptional, because the total *virF* mRNA signal does not change between the wild-type and ΔRS *S. flexneri* strains (Fig. 7C).

The ΔRS bacteria are longer than wild-type *S. flexneri* when grown *in vitro* and demonstrate extensive cell elongation within Henle cells (Fig. 8A). Longer length of ΔRS cells has been demonstrated in *E. coli*, and this was found to be a direct effect of reduced levels of (p)ppGpp, rather than an indirect effect due to changes in growth rate (Büke et al., 2022). The elongation of *S. flexneri* ΔRS cells is recapitulated when the bacteria are grown in LB medium containing increased NaCl or in medium with restricted iron availability, whereas wild-type cells do not change their length in response to these stressors (Fig. 8B and 8C). These data suggest that the ΔRS strain is exposed to increased osmotic or ionic pressure, low iron concentration, or other environments inside the Henle cells that lead to a stress-related elongation phenotype. Exposure to osmotic stress inside the host cells is consistent with data showing that the *proVWX* genes encoding the osmoregulatory ProU system proteins are upregulated during *S. flexneri* infection of HeLa cells and macrophages (Lucchini et al., 2005). Further, mutants of *proV* in *S. sonnei* are defective for intracellular growth (Mahmoud et al., 2017). An RNA-seq experiment profiling genes in *E. coli* that are modulated by (p)ppGpp shows that *proV* is activated three-fold by (p)ppGpp (Sanchez-Vazquez et al., 2019). Thus, one role of (p)ppGpp *in vivo* may be to induce expression of *Shigella* osmoregulation genes or other genes for adaptation to stress when the bacteria are growing in the host cell cytosol.

Our results indicate that SpoT can generate the levels of (p)ppGpp that are sufficient for *Shigella* virulence. This adds to the list of pathogens that use this alarmone to modulate metabolic pathways and virulence factors within the host. Although SpoT generates basal levels of (p)ppGpp, in the case of intracellular *S. flexneri,* our results suggest that SpoT is necessary to generate elevated levels of (p)ppGpp in response to nutrient limitation and/or physiological stress. The ability of (p)ppGpp to enhance virulence appears to be linked to the maintenance of VirF levels; however, the precise molecular mechanism through which (p)ppGpp acts in *S. flexneri* remain to be determined.

## MATERIALS AND METHODS

### Strains, Media, and Growth Conditions

Strains are listed in Table 1. All strains were maintained at -80°C in Tryptic Soy Broth (TSB) containing 20% (v/v) glycerol. *S. flexneri* strains were grown on TSB agar (TSB, 1.5% agar [w/v]) containing Congo red dye (0.01% [w/v]; Sigma) at 37°C, and red (i.e., putative virulent) colonies were selected (Payne & Finkelstein, 1977). Isolated colonies were inoculated in LB broth and grown at 30°C with shaking for 16-18 h, and then sub-cultured 1:100 (wild-type) or 1:50 (ΔRS) and grown at 37°C for assays. Antibiotics were used at the following concentrations: chloramphenicol, 25 μg/ml; ampicillin, 25 μg/ml; and kanamycin, 50 μg/ml. X-gal (0.2 mg/ml) was added to L-agar to determine β-galactosidase production.

**Table 1.** Strains and plasmids.

| Strain or plasmid name: | Description | Reference |
| --- | --- | --- |
| ***E. coli*:** |  |  |
| TOP 10 | Cloning strain | Thermo Fisher |
| ***S. flexneri* 2a:** |  |  |
| Wild type | 2457T wild type, serotype 2a (AE014073.1) | Walter Reed Army Institute of Research |
| $\Delta R$ | 2457T $\Delta relA::cam$ | This study |
| $\Delta RS$ | 2457T $\Delta relA \Delta spoT::cam$ | This study |
| *icsA* or $\Delta icsA$ | 2457T *icsA::kan* | H. Marman, R. Rossi, Payne lab |
| CFS100 | *S. flexneri* 2457T cured of virulence plasmid | C. Fisher, Payne lab |
| **Plasmids:** |  |  |
| pKD3 | Cam cassette template | (Datsenko & Wanner, 2000) |
| pKD46 | Ts Red Recombinase | (Datsenko & Wanner, 2000) |
| pCP20 | Ts FLP recombinase | (Datsenko & Wanner, 2000) |
| pWKS30 | Low-copy number vector | (Rong Fu Wang & Kushner, 1991) |
| $\Delta RS/pWKS30-spoT$ | *spoT* in pWKS30 | This study |
| pBAD24 | Arabinose inducible protein expression vector | ATCC. #87399 |
| $\Delta RS/pBAD24-spoT$ | *spoT* in pBAD24 | This study |
| pQF50 | Promoterless *lacZ* expression vector | (Farinha & Kropinski, 1990) |
| *csgD*-pQF50 | *csgD*-5'UTR region cloned upstream of *lacZ* | This study |
| virF-S-tag | pWKS30 plasmid containing a C-terminal S-tag | (Xerri & Payne, 2022) |
| AKS2000 | *hfq* in IPTG inducible vector | (Sharma & Payne, 2006) |
| pmCherry | Expression vector that encodes mCherry | Clontech/Takara cat # 632522 |

### Construction of bacterial mutant strains

Both *S. flexneri relA* (ΔR, single deletion) and *relA spoT* (ΔRS, double deletion) mutants were constructed using λ-Red-mediated recombination (Datsenko & Wanner, 2000). The *relA* mutant was created first and formed the background for the *relA spoT* double mutant strain. A PCR product was generated by amplifying the chloramphenicol resistance cassette from pKD3 with primers (Table 2) containing overhangs complementary to sequences flanking the gene to be mutated. The amplicon size was verified via agarose gel electrophoresis and introduced by electroporation into *S. flexneri 2457T* expressing λ-Red-recombinase from plasmid pKD46 (Datsenko & Wanner, 2000). Recombinants were purified first on plates without antibiotic then on plates with chloramphenicol. Correct replacement of the chromosomal gene with the chloramphenicol resistance cassette was verified via PCR and DNA sequencing.

**Table 2.**
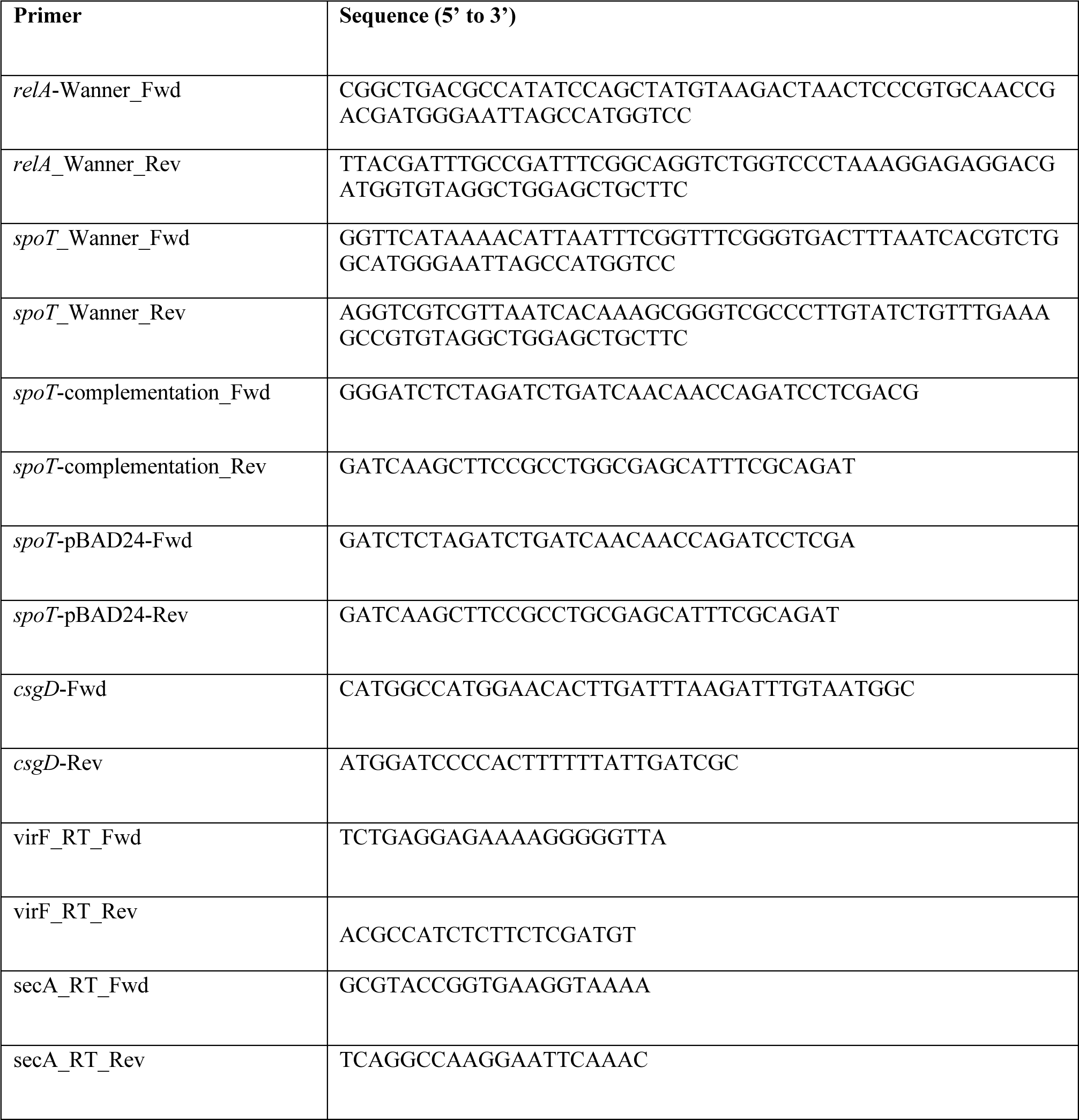
Primers.

### Plasmid construction

pSpoT-pWKS30: A SpoT expression plasmids was constructed by amplifying *spoT* from *S. flexneri* 2457T and ligating it into the SmaI site of pWKS30. Integration into the plasmid was screened initially via blue-white screening, and then confirmed via DNA sequencing.

pSpoT-pBAD-24: A plasmid expressing *spoT* under an inducible arabinose promoter was generated by amplifying *spoT* from *S. flexneri* with primers that contained the restriction enzyme sites XbaI and HindIII. This construct was inserted into pBAD-24 and the sequence was verified by Sanger sequencing.

pCsgD-pQF50: A plasmid containing the (p)ppGpp-regulated promoter of *csgD* fused to *lacZ* was generated as follows. The *csgD* promoter (195 bp) was amplified from *S. flexneri* 2457T with primers *csgD*-Fwd and *csgD*-Rev, and size-verified by agarose gel electrophoresis. This sequence is identical to that reported for *E. coli* (Sanchez-Vazquez et al., 2019). The fragment was digested with Nco1 and BamHI, and ligated into a similarly digested *lacZ* fusion plasmid pQF50 (Farinha & Kropinski, 1990). Insertion was initially verified by blue-white screening and confirmed by DNA sequencing.

### Cell Culture Assays

#### Invasion Assay

Bacteria were grown to mid-log phase (OD_650_ = 0.5-0.8) and 10^8^ cfu were added to a confluent monolayer of Henle cells in 35 mm, 6-well, polystyrene plates (Corning) and centrifuged onto the monolayer at room temperature for 10 min at 1,000 x g. Plates were incubated for 30 min, then washed four times with PBS-D (26.5mM KCl, 1.5mM K_2_HPO_4_, 137mM NaCl, 8mM Na_2_HPO_4_). Minimal Essential Medium (MEM) containing gentamycin (20 µg/ml) was added, and the plates were incubated for an additional 60 min. The monolayers were washed four times with PBS-D, 1ml of warmed 5% saponin (w/v in PBS) was added, and the plates were incubated for 10 min at 37 °C to lyse the Henle cells. The lysate was serially diluted and plated on TSB. The input was also determined by serially dilution and plating. Each experimental condition was conducted at least three times.

#### Plaque assay

Plaque assays were a modification of the protocol of Oaks et al. (Oaks et al., 1985) Assays were conducted as for the invasion assay, above, with the following modifications: 10^4^ cfu of *S. flexneri* were added to confluent monolayers, and the plates incubated for a total of 72 hours, with a change of medium after 24 h incubation. Monolayers were stained with either Wright-Giemsa stain (Camco) for visualization or fixed with 80% methanol for 5 minutes and then stained with 0.5% crystal violet in 25% methanol. Where indicated, arabinose (0.4%) or IPTG were added to induce expression of cloned genes.

### Detection of IcsA

#### In vitro

To determine surface localization of IcsA, bacteria were grown to mid-late log phase, centrifuged at 3,500 x g for 5min at room temperature, and resuspended in 4% paraformaldehyde (PFA) for 10-20 min. Cells were washed three times in saline (0.9% NaCl), and then spotted onto glass slides. Primary antibody (rabbit #35, Edwin Oaks, Walter Reed Army Institute of Research) was diluted 1:120 in PBS +2.5% BSA, spotted onto the cells, and incubated for 1 h at room temperature. The slides were washed three times with PBS, and 1:100 dilution of fluorescent isothiocyanate (FITC)-conjugated goat anti-rabbit secondary (Invitrogen, F2765) antibody diluted in PBS + 2.5% BSA was added for 2 hours at 4°C. The slides containing the cells were washed three times in PBS, and mounting media and a coverslip applied. Samples were cured overnight using Prolong Diamond (Thermo Fisher P36965) in the dark and imaged the next day. Where indicated, cephalexin (64 µg/ml) was added to induce elongation (Chung et al., 2011) prior to fixing and staining the cells.

To image both the cytoplasmic and secreted IcsA populations in the bacterial cells, the method above was a modification of the protocol of Campbell-Valois et al. (Campbell-Valois et al., 2014). Briefly, after fixing, 10 µl of bacteria were spotted onto glass coverslips and dried, then 0.1% Triton was added, and incubated for 5 min. The coverslip was washed carefully and dried at room temperature. Lysozyme solution (50 mM glucose, 5 mM EDTA, 5 mg/ml lysozyme in PBS) was applied onto the bacterial spot for 25 min at 37°C, then washed three times with PBS. Anti-IcsA primary antibody (rabbit #35, Edwin Oaks, Walter Reed Army Institute of Research) was applied for 1h at RT, washed three times with PBS, followed by fluorescent isothiocyanate (FITC)-conjugated goat anti-rabbit secondary antibody for 1h. The coverslips were gently washed three times with PBS and Prolong Diamond (Thermo Fisher P36965) mounting medium was applied and the coverslip cured overnight at room temperature until imaging.

#### In vivo

IcsA on intracellular bacteria was visualized as previously described (Purdy et al., 2007). Briefly, bacteria were grown to mid-log phase, centrifuged at 3,400 x g for 5 min at room temperature, and resuspended in saline solution (0.9% NaCl). These bacteria were then applied onto 70% confluent Henle-407 cells at ∼2x10^7^ cfu/ml. The plates were centrifuged at (1000 x g) for 10 min at room temperature and placed in a 37°C incubator with 5% CO_2_ for 30 min. Plates were washed with PBS and MEM containing gentamicin was applied onto the cells. The cells were incubated for 3h. Cells were fixed with 4% paraformaldehyde (PFA) for 10 min, permeabilized with 0.1% Triton X-100 for 10 min, and blocked with 5% BSA for 30 min. Cells were labeled by indirect immunofluorescence overnight using rabbit polyclonal antibody against IcsA (Rabbit 35) provided by Edwin Oaks (Walter Reed Army Institute of Research) diluted 1:100 in PBS and 2.5% BSA, and a fluorescent isothiocyanate (FITC)-conjugated goat anti-rabbit secondary antibody diluted 1:100.

#### Microscopy

Fluorescence microscopy was performed at the Center for Biomedical Research Support Microscopy and Imaging Facility at UT Austin (RRID# SCR_021756). Images were acquired using a Nikon W1 spinning disk confocal microscope containing a Yokogawa W1 scanning head with multiple pinholes for ultrafast confocal imaging, using the 488 and 561 nm excitation lasers and the Plan Apochromat λ 100x (Oil); NA - 1.45; WD - 0.13 mm objective. Images were range-adjusted in ImageJ. Micrographs were visualized via Fiji/Image J (Schindelin et al., 2012), and capped cells were reported as a percentage of total cells (% *in vitro* or *in vivo* capped *S. flexneri*).

#### Western Blotting Assays

Bacterial proteins were separated on 10% or 15% (for Hfq detection) polyacrylamide SDS gels. Post-electrophoresis, proteins were transferred to a 0.45 μm pore size nitrocellulose membrane (GE Healthcare) and incubated with rabbit polyclonal anti-Hfq antibody (Andrew Feig, Indiana University, Bloomington, Indiana), monkey *S. flexneri* convalescent antiserum (Edwin Oaks, Walter Reed Army Institute of Research), or anti S-tag (MA1981, Life Technologies). Proteins were detected using horseradish peroxidase (HRP)-conjugated goat anti-rabbit, anti-mouse, or anti-human antibody (all diluted 1:10,000). Signal was detected by developing the blot with a Pierce ECL detection Kit (Thermo Fisher Scientific).

#### Western Blot Quantitation

Quantitation of western blots were conducted using Fiji. Specifically, a rectangle was drawn to encompass all the bands representing the virulence protein profile per sample. The rectangle was propagated to all the lanes on the gel image, and using the “Plot-lane” function, isolated peaks corresponding to each band intensity was generated. A baseline for each set of peaks was drawn using the “line” tool, and using the “Wand-tracing” tool, the areas under the peak were quantified. The background signal was subtracted from the signal generated from the “CFS100” lane. The resultant values corresponding to each band intensity were normalized to the wild-type and reported as a fraction of the wild-type signal. Three biological replicates were analyzed. Three biological replicates were analyzed, and significance tested using the Students *t* test, p less than 0.05.

#### Miller Assay

Miller assay to quantitate β-galactosidase activity was conducted using the modified Miller assay as described by Zhang and Bremer (1995), using the protocol available on openwetware.org (https://openwetware.org/wiki/Beta-Galactosidase_Assay_(A_better_Miller).

#### Genomic alignments

Alignments of *relA* and *spoT* were performed using Rapid Annotation using Subsystem Technology (RAST) (Aziz et al., 2008; Brettin et al., 2015; Overbeek et al., 2014). Source genomic DNA was acquired from NCBI using the following accession numbers: AE014073.1 (*S. flexneri*), NZ_CP026795.1, (*S. boydii* ATCC BAA-1247), NZ_CP055292.1 (*S. sonnei* SE6-1), NZ_CP055055.1 (*S. dysenteriae*), and NC_002695.2 *(E. coli* 0157:H7 str. Sakai).

#### qPCR

*S. flexneri* strains were grown in LB to late log phase. Four milliliters of the cell suspension were mixed with 1 mL of an ice-cold solution containing 95% ethanol and 5% phenol (pH 4.5) and chilled on ice. The cells were pelleted by centrifugation, resuspended in 100 μL of lysozyme (1 mg/mL) in Tris-EDTA (TE) buffer, and incubated at room temperature for 5 min. One ml of RNA-Bee (Tel-Test Inc.) and 200 μL of chloroform were added, and the solution was centrifuged at 4°C and 21,400 × g for 15 min. The aqueous phase was collected, mixed with an equal volume of isopropanol, and stored at −80°C overnight. Samples were centrifuged at 21,400 × g for 20 min to pellet precipitated RNA, which was washed once with ice-cold 75% ethanol, air dried, resuspended in water, and DNase treated (Turbo DNA-free; Invitrogen). Two micrograms of RNA harvested from the wild-type and ΔRS strains were reverse transcribed to generate cDNA using the Superscript III kit (Thermo Fisher). VirF primers (Xerri & Payne, 2022) for quantitative PCR (qPCR) were designed using Primer3 (http://bioinfo.ut.ee/primer3-0.4.0/). For real-time qPCR, cDNA was diluted 1:10 and mixed with Power SYBR green (Thermo Fisher). The qPCR was run on an Applied Biosystems ViiA7 instrument as in (Butz et al., 2019) Relative expression of virulence genes was calculated by using the threshold cycles (ΔΔCT) method and normalized to *secA*.

#### Bacterial culture with NaCl or iron chelator

Bacterial cultures were inoculated from single colonies of either the wild-type or ΔRS strain and grown overnight at 30°C. The cultures were then subcultured 1:100 for the wild-type and 1:50 for the ΔRS and grown to mid-log phase, and then subcultured once more in LB, or LB containing either 340mM NaCl or 250µg/ml of the iron chelator ethylenediamine di(*ortho*-hydroxyphenylacetic acid) (EDDA) (Wyckoff & Payne, 2011). Cells were grown for 3 hours at 30°C in the presence of the stressing agents, and then 1ml of culture harvested for fixing. Bacterial cells were centrifuged at 3.4K x g for 5 minutes at room temperature and resuspended in 4% PFA for 10 minutes. The cells were then washed once with PBS, pelleted, and resuspended in 200-300µl PBS. Five to seven microliters of the cell suspensions were spread onto glass slides and dried at room temperature. A coverslip containing Prolong Diamond (Thermo P36965) mounting media was applied onto the dried smears and samples were cured overnight. Bacteria were imaged using phase contrast microscopy using a 100x objective, and lengths were quantified using Fiji/ImageJ. One hundred cells were measured from each condition, from three biological replicates per experiment. The lengths of all the cells are reported (Fig. 8).

## Acknowledgements

Research reported in this publication was supported by the National Institute of Allergy and Infectious Disease of the National Institutes of Health under award number R37NIAID16935. We thank Alexandra Mey, Camilo Gómez-Garzón, and Carolyn Fisher for helpful discussions and reagents.

